# A Role for Maternal Factors in Suppressing Cytoplasmic Incompatibility

**DOI:** 10.1101/2020.06.27.174805

**Authors:** AJM Zehadee Momtaz, Abraham D. Ahumada Sabagh, Julian Gonzalez Amortegui, Samuel A. Salazar, Andrea Finessi, Jethel Hernandez, Steen Christensen, Laura R. Serbus

## Abstract

*Wolbachia* are maternally transmitted bacterial endosymbionts, carried by approximately half of all insect species. *Wolbachia* prevalence in nature stems from manipulation of host reproduction to favor the success of infected females. The best known reproductive modification induced by *Wolbachia* is referred to as sperm-egg Cytoplasmic Incompatibility (CI). In CI, the sperm of *Wolbachia*-infected males cause embryonic lethality, attributed to paternal chromatin segregation defects during early mitotic divisions. Remarkably, the embryos of *Wolbachia-*infected females “rescue” CI lethality, yielding egg hatch rates equivalent to uninfected female crosses. Several models have been discussed as the basis for Rescue, but none have been demonstrated in robust CI models, which are genetically intractable. As such, the extent of host involvement in Rescue remains untested. In this study, we used a chemical feeding approach to assess maternal contributions to CI suppression in *Drosophila simulans*. We found that uninfected females exhibited significantly higher CI egg hatch rates in response to seven chemical treatments that affect DNA integrity, cell cycle control, and protein turnover. Three of these treatments suppressed CI induced by endogenous *w*Ri *Wolbachia*, as well as an ectopic *w*Mel *Wolbachia* infection. When framed in terms of existing literature, the results implicate DNA integrity as a dynamic aspect of CI suppression for different *Wolbachia* strains. The framework presented here, applied to diverse CI models, will further enrich our understanding of host reproductive manipulation by insect endosymbionts.

## INTRODUCTION

Endosymbiosis is a specialized form of interaction, with one organism dwelling inside the cells and tissues of another (1). The bacterium *Wolbachia pipientis* is one of the most widespread endosymbionts, carried by half or more of all insect species (2, 3). *Wolbachia* are gram negative bacteria that belong to the alpha-protobacterial class *Rickettsiales. Wolbachia* are maternally transmitted, with the efficacy of transmission dependent on the bacteria being loaded into eggs (4-12). *Wolbachia* commonly modify host reproduction to favor the success of infected females. This is accomplished by induction of parthenogenesis, male killing, feminization and sperm-egg cytoplasmic incompatibility (13-18). Though *Wolbachia* interactions with their host appear generally commensal, this extent of host manipulation classifies *Wolbachia* bacteria as reproductive parasites.

Cytoplasmic incompatibility (CI) is the most widely known of all *Wolbachia*-induced reproductive manipulations (4, 19, 20). CI is characterized by embryonic lethality in crosses between uninfected females and *Wolbachia*-infected males (13, 21-28). By contrast, *Wolbachia*-infected females are compatible with both uninfected and *Wolbachia*-infected males, with viable progeny produced by both types of crosses. The ability of embryos from *Wolbachia*-infected females to survive the *Wolbachia*-modified sperm is known as “Rescue.” Compatibility is conferred by specific pairings of sperm modification (mod) and rescue capacity (resc) associated with different *Wolbachia* strains (20, 29-35). With infected females favored by elimination of incompatible embryos, the CI/Rescue paradigm effectively drives host population replacement in natural populations as well as in applied, vector management scenarios (36-43).

The cellular basis of CI has been a point of interest for many years. Cytological experiments indicate that mitotic defects are a consensus feature of CI across *Wolbachia*-host systems. Specifically, paternal chromatin remains at the metaphase plate while maternal chromatin segregates to opposite poles in anaphase, resulting in chromosome bridging and aneuploidy (44-51). Studies from *Nasonia* and *Drosophila simulans* have suggested these mitotic defects are produced from a timing mismatch between male and female pronuclei at the first mitotic division, which must be reconciled in order to enable Rescue (46, 48, 49). A separate line of work implicated *Wolbachia-* induced oxidative damage to spermatocyte DNA as a contributor to CI lethality (52) with the implication that DNA damage prevention and/or repair in the embryo is important in conferring Rescue. This model is consistent with a body of literature on *Wolbachia* and induction of oxidative stress (53-59).

The most recently presented model is that *Wolbachia* CI factors (Cifs) are responsible for both CI and Rescue (60-62). Bioinformatic predictions, and studies using yeast and *Drosophila melanogaster* models, have indicated that CifB proteins have either deubiquitlase activity (CidB), nuclease activity (CinB), or both (CndB) (62-66). CifB is required to induce CI (50, 60, 65, 67), possibly through direct impacts on sperm chromatin remodeling during spermatogenesis (66). By contrast, CifA can Rescue classical CI phenotypes induced by *Wolbachia* (61), as well as CI associated with dual expression of CifA and CifB in transgenic males (65, 68).

Despite these advances, the lack of a genetically tractable system for analyzing robust CI and Rescue phenotypes has hindered mechanistic understanding. Recent studies of Cif function in the *D. melanogaster* model have provided unprecedented insight, with the intrinsic caveat that *D. melanogaster* is a mild CI system. CI lethality is induced by newly eclosed males, particularly when the first-emerging males of a population are used for the CI cross, but this CI capacity diminishes rapidly with male age (60, 69, 70). The transience of CI appears attributable to the host background, as transfer of *w*Ri *Wolbachia* into *D. melanogaster* also elicits a mild CI response (71), unlike the robust CI expression associated with the *D. simulans* host (27, 42, 72, 73). To date, the mechanism of Rescue has not been functionally demonstrated in a robust CI model.

A long-standing question has been to what extent embryonic factors contribute to the mechanism of Rescue. If Cif proteins act exclusively in terms of a toxin-antitoxin system, with CifA suppressing CifB function via direct binding (64), no maternal involvement is required for Rescue. Consistent with this model, direct binding has been demonstrated for multiple cognate pairs of CifA and CifB proteins (62, 65, 66). However, *Wolbachia* gene expression patterns also indicate a natural temporal disconnect between Cif proteins during *D. melanogaster* development (63). CifB reportedly shows high expression in early-to mid-embryogenesis, whereas CifA is more highly expressed in late embryogenesis (63). This observation leaves open the possibility of additional roles for Cif proteins, or other as-yet-unidentified factors, in conferring Rescue ability upon embryos. A detailed analysis did not identify contributions by additional *Wolbachia-*generated factors (74), thus a role for maternal contributions remains to be investigated.

To increase our understanding of Rescue, we tested the hypothesis that Rescue is supported by endogenous cellular mechanisms of insects. If maternal factors contribute to Rescue, one prediction would be that modifying these same pathways should confer CI suppression abilities upon uninfected embryos. To test this, we applied a chemical feeding approach to alter candidate cellular processes in the classical *Drosophila simulans* CI/Rescue model. After developing and optimizing a plate-based screening assay, 24 candidate chemicals were tested that address existing mechanistic hypotheses about CI and Rescue. CI induced by the endogenous *w*Ri *Wolbachia* strain, as well as a transinfected *w*Mel strain, were investigated. Analyses of egg hatch were performed to determine the extent of chemical impact on CI, with results evaluated in light of existing CI/Rescue models, as described below.

## MATERIALS AND METHODS

### Fly stocks and rearing conditions

The *w*Ri *Wolbachia* strain, endogenous to *Drosophila simulans*, used in this study was originally described by Hoffman, Turrelli, and Simmons (27). The uninfected *D. simulans* strain (w^-^) is of the same genetic background, as it was this original line cured of *Wolbachia* with tetracycline. The *w*Mel trans-infected line was created with *Wolbachia* from *D. melanogaster* (73), backcrossed into the cured fly stock for six generations to standardize the *D. simulans* genetic background. We previously confirmed the identity of *w*Ri and *w*Mel in *D. simulans*, and verified that the *w*Mel transinfection matches the standard *w*Mel strain carried by most *D. melanogaster* stocks (75).

All the flies used in this study were maintained at 25°C on a 12h light/dark cycle using an Invictus Drosophila incubator (Genessee Scientific, USA). Flies were raised in standard 6oz square bottom polyethylene bottles containing 25-30ml of fly food (described later). Each stock bottle was seeded by approximately 80-100 flies, a mixture of both male and female, and incubated for 3-6 days. After this period, flies were either transferred to a new bottle or discarded. Flies used to seed all bottles were discarded by 12-15 days of age. To collect virgin flies, stock bottles were completely cleared, and rechecked by eye to verify the absence of flies. Newly eclosed flies were collected 5-8 hours later using standard CO2 gas pads. Males and females were separated and temporarily stored in narrow polypropylene vials until loading into treatment vials/plates, as described below. To avoid damage from prolonged exposure to CO2, fly sorting was limited to a 20-25 min time frame.

### Microbial 16S rRNA gene sequencing

To determine the infection status in *D. simulans* flies, microbial 16S rRNA gene sequencing was performed. Ovaries were dissected from uninfected, *w*Ri-infected and *w*Mel-*D. simulans* females, followed by 16s rRNA gene sequencing carried out as previously described. (76). Each sample represents ovarian content from 20 flies.

### Food preparation

The stock food used in this study was made as per a standard Bloomington Stock Center recipe, described earlier (https://bdsc.indiana.edu/information/recipes/bloomfood.html; (77). For chemical feeding assays, concentrated stock solutions of each chemical were prepared using an appropriate solvent, dependent on necessary final concentration. To make chemical food for independent experiments, the appropriate amount of stock solution was mixed with melted food and mixed thoroughly by stirring. This food was transferred to either vials or plate wells immediately, before cooling and solidifying. The same amount of solvent was mixed with standard food to create a parallel “control food” condition in each experiment. Vial-based trials contained 3-5ml of food per vial, while in the 24-well plate format, each well contained 800 μL of food.

### Dose response curve preparation

To determine the appropriate feeding concentration for each chemical, a range of 6-7 doses was empirically tested for each compound. The range of concentrations used was based on information available in existing literature. Each chemical was diluted to the appropriate concentration in beakers containing 10 mL standard food with Brilliant Blue G food added (Acros Organics). All content was mixed thoroughly for 30 seconds, then divided equally into 2 treatment vials and cooled under the fume hood for ∼2 hours to prevent condensation from colleting along the sides of the vials. In each case, one treatment vial was immediately used, and the second vial was plugged with rayon, wrapped in aluminum foil, and stored in a sealed container at 4°C for use 6 days later.

To carry out dose-response testing, 6 uninfected male and 6 uninfected female *D. simulans* flies were incubated in the first set of treatment vials. On the 6^th^ day, the flies were transferred to the second corresponding treatment vials for an additional 6 days. Adult mortality, egg lay, egg hatch and larval development were qualitatively scored for treatment vials and corresponding controls across the 12-day period. All measurements were scored by comparing the treatment vials with the control vials. If treatment resulted in some negative effects during later days (7-12) compared to control, or if treatments showed consistent developmental defects across the 12-day span. These were scored respectively as “+”, “some” or “-“. The highest concentration of chemical with no adverse effect on flies as per the above criteria was selected for subsequent feeding assays. Dose response curves for dual drug treatment combinations were carried out similarly. Two independent biological replicates were performed for all dose-response experiments.

### CI suppression tests

#### In the vial-based format

Virgin *D. simulans* flies were kept in vials containing 3-5ml of food and incubated for 3 days. Females were split between treatment food and control food conditions, whereas infected males were exposed to standard food only. In all cases, flies were grouped, with 15-20 flies per vial. On day 3, male and female flies were combined and transferred to fresh vials of standard food for an 8-hour mating period. Depending upon the experiment, this was done as single pair matings or as mass matings of 30-40 flies, using equal numbers of males and females. Afterwards, male flies were discarded and female flies were transferred to their original treatment vials. At day 4, individual females were split up into separate vials that sustained their existing treatment conditions, with the addition of blue food coloring to the food to improve egg visibility. At day 5, the female flies were discarded, and the vials were incubated an additional 24 hours to allow for egg hatch. At day six, all the unhatched and hatched eggs were counted to determine the egg hatch rate. Each treatment was run as two separate biological replicates.

#### In the plate assay format

Corning 24-well plates (Cat# 3738) were set up with 16 wells carrying control food and 8 wells carrying treatment food. Each well carried a total volume of 800μL. 10 *D. simulans* virgin females were added to each well and incubated for 3 days. Uninfected females were added to 8 standard food wells for use as the CI control, and to 8 treatment wells to test for chemical suppression of CI. Infected females were added to 8 standard food wells, for use as a Rescue control. Infected males were incubated separately on standard food for 3 days, at a density of 45 flies or fewer per vial. At day 3, the females were transferred to a new plate carrying standard food only. 10 infected male flies were added to each well for an 8-hour mating. Afterwards, males were removed, and female flies returned to their respective wells in the original treatment plate. At day 4, females were transferred to a fresh plate that contained the same treatment condition per well, now including blue food coloring dye. At day 5, the female flies were discarded. At day 6, egg hatch was scored per well, yielding an egg hatch rate for the fly population of each well. For further details on plate management and assay execution see Additional file 1. Two or more independent biological replicates were performed for each plate assay experiment. For rigor and consistency across the dataset, only plates that showed a 12% or lower hatch rate for the CI control were scored.

### Statistical analysis

Chi tests of goodness of fit were performed manually as per standard procedures (78). A Bonferonni correction was applied, so that alpha values were scaled to the number of data categories analyzed (78). Z’ values were calculated as previously (79, 80). As previously, the IBM SPSS v.23 analysis package was used for all other statistical test (76, 81). The data were analyzed for normality using the Shapiro-Wilk test and for homogeneity of variance by Levene’s test (82-84). For the data with normal distribution, mean differences were evaluated using a T-test if variance was homogeneous, and Welch’s T-test if it was not (78, 85). If the data did not fit the normal distribution, the Mann-Whitney U test was performed for data showing homogeneity of variance, and an independent T-test was performed with bootstrapping as an approximation when variance was uneven (85, 86). To assess the power of different sample sizes, we also used a bootstrap procedure in MATLAB™ (Mathworks, Natick MA) that randomly sub-samples from the data to determine the sample size required to meet specified *p*-values (76).

## RESULTS

### Verification of *D. simulans* endosymbiont identity by 16S analysis

The *Drosophila simulans* flies used in this study carry the *Wolbachia* strain *w*Ri as a natural infection (27), or *w*Mel as a transinfected strain (73). Female flies from these strains have been shown by DNA staining to exhibit nucleoids in their germline cells that are consistent with *Wolbachia* infection (75, 87). These lines have also been confirmed as PCR-positive for the *Wolbachia surface protein* gene, and the *Wolbachia* strain identities have been confirmed by sequencing (75). Use of these detection methods, while consistent with expected *Wolbachia* identities, does not rule out the possible presence of other bacterial endosymbionts.

To independently confirm the identity of the germline bacteria carried by *Wolbachia*-infected flies, 16S rDNA microbiome analyses were carried out as previously described (76). Ovary tissue samples were analyzed from both uninfected and *Wolbachia*-infected *D. simulans* lines. The data indicate *Wolbachia* spp. as the predominant taxon carried by both *w*Ri- and *w*Mel-infected tissues, with 94.5-98.3% of the reads representing the *Wolbachia* genus (Figure 1) (Table S1) (Additional File 2). The other non-*Wolbachia* taxa detected in the *Wolbachia-*infected ovary samples paralleled that of the uninfected control (Figure 1). As this is a non-sterile assay, these signatures likely reflect the microbiome of the cuticle, the body cavity and residual contamination of dissection equipment (76). Notably, the *Wolbachia-*infected samples show no evidence of other *Drosophila-*resident symbionts such as *Spiroplasma*. Thus, *Wolbachia* endosymbionts represent the vast majority, if not all, of the microbiome carried by maternal germline cells that generate Rescue-capable eggs.

**Figure 1.**
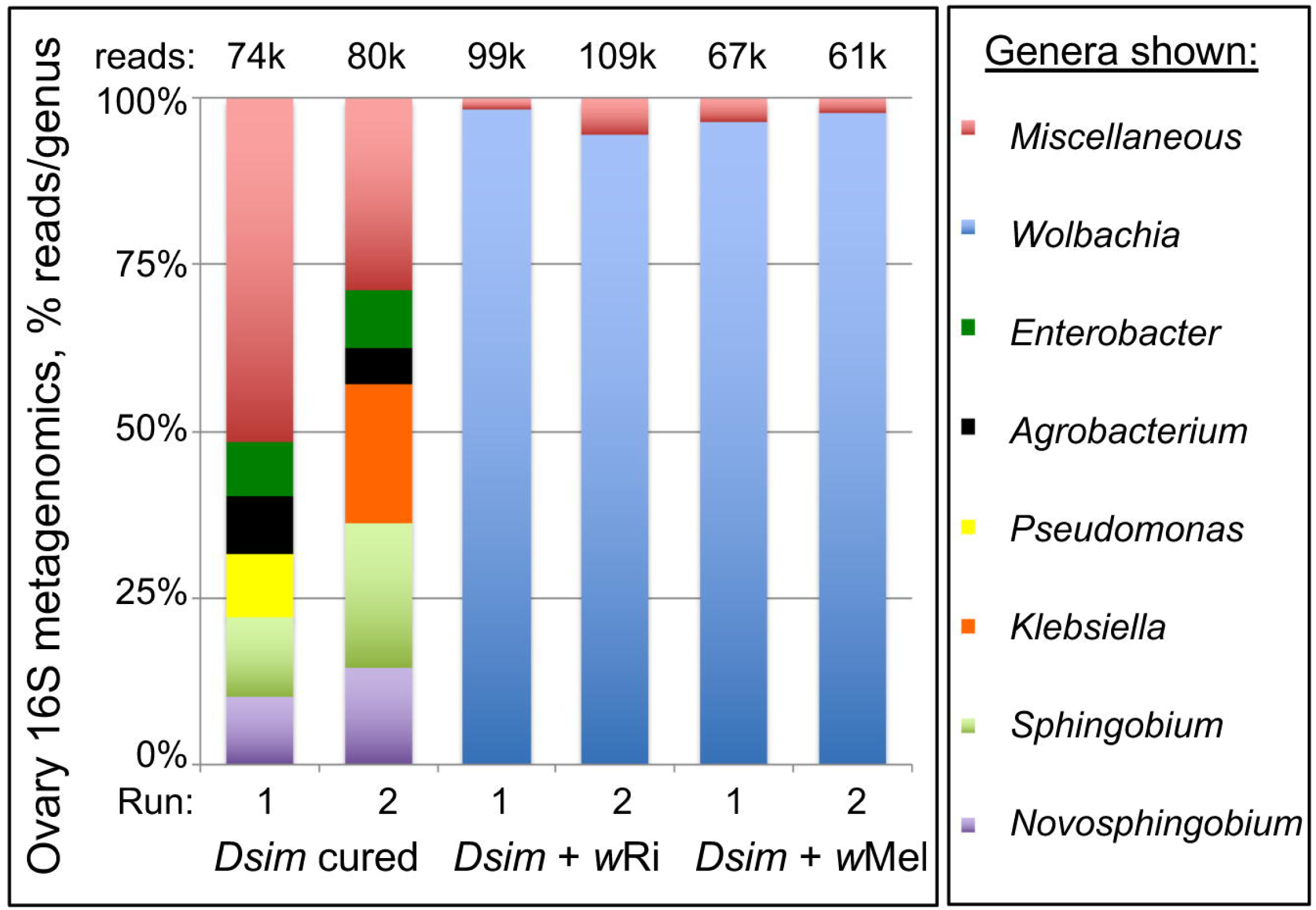
16S microbiome profiles associated with *D. simulans* ovary tissues. Uninfected and *Wolbachia*-infected tissues are shown. Top five most abundant genera that equal or exceed 1% abundance per sample are shown. For further details, see Table S1 and Additional File 2.

### Confirmation of baseline CI and Rescue phenotypes from *D. simulans*

Previous studies have demonstrated that natural (*w*Ri) as well as transinfected (*w*Mel) *Wolbachia* strains induce robust CI and Rescue effects in *D. simulans* (27, 28, 73). To determine the strength of CI and Rescue in current laboratory settings, group mating assays were performed and egg hatch outcomes were scored for individual females. Crosses of infected females to infected males, referred to as the Rescue, yielded a 92% hatch rate. This was not significantly different from the uninfected Control cross (*p* > 0.05; adjusted *α* = 0.0083) (Table 1). By contrast, the CI egg hatch rates ranged from 4-10% for *w*Mel and *w*Ri. This represents a significant reduction in hatch rates, as compared to both Control and Rescue crosses (*p* < 0.001; adjusted *α* = 0.0083) (Table 1). This outcome is consistent with the expectation of strong CI and Rescue phenotypes associated with *D. simulans*.

**Table 1.**
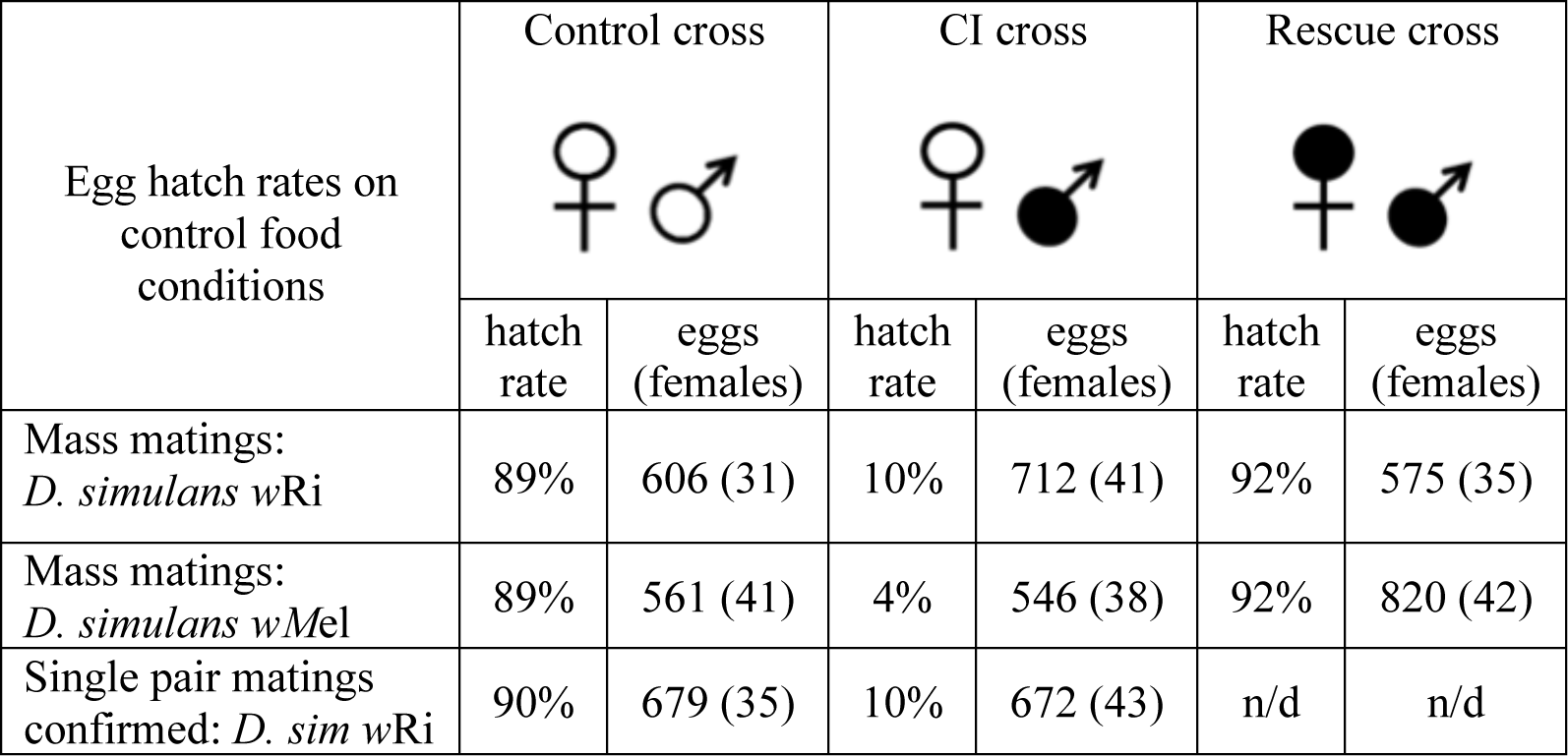
Egg hatch rates from CI-related crosses on control food

| Egg hatch rates on control food conditions | Control cross |  | CI cross |  | Rescue cross |  |
| --- | --- | --- | --- | --- | --- | --- |
|  | hatch rate | eggs (females) | hatch rate | eggs (females) | hatch rate | eggs (females) |
| Mass matings:<br>*D. simulans* wRi | 89% | 606 (31) | 10% | 712 (41) | 92% | 575 (35) |
| Mass matings:<br>*D. simulans* wMel | 89% | 561 (41) | 4% | 546 (38) | 92% | 820 (42) |
| Single pair matings confirmed: *D. sim* wRi | 90% | 679 (35) | 10% | 672 (43) | n/d | n/d |

*Wolbachia* infection has been reported to alter pheromone release and perception by other species of *Drosophila*, leading to altered mating patterns (88, 89). Thus, it is formally possible that low egg hatch in incompatible crosses, normally attributed to *w*Ri-induced CI lethality, instead reflects a failure to mate. To distinguish between these possibilities, single pair matings were set up, with egg hatch scored only for vials in which mating was visually confirmed. In these experiments, hatch rates remained low for CI crosses (10%) as compared to Control crosses (90%) (Table 1) (*p* < 0.001; adjusted *α* = 0.0125). The results of these mating-confirmed crosses closely parallels that shown above for group matings. This demonstrates that the low hatch rates currently associated with *w*Ri-induced CI are attributable to the reduced viability of eggs laid by uninfected *D. simulans* females.

### Demonstrating use of small molecule inhibitors to suppress CI phenotypes

CI embryos have previously been shown to exhibit defective incorporation of maternal histones into paternal DNA (49). We reasoned that chromatin-modifying compounds may also be able to confer CI suppression in a manner analogous to natural Rescue. It is known that acetylation of histones lowers their affinity for DNA and loosens chromatin structure, whereas removal of acetyl groups by histone de-acetylase (HDAC) enzymes reverses this effect (90). Since HDAC inhibitor compounds are commonly used in animal models and clinical settings (90-94), an array of well-established compounds is available for testing. Thus, we established a chemical feeding protocol to test a role for chromatin remodeling in CI suppression.

One of the most well-known HDAC inhibitors is the non-toxic, short chain fatty acid butyric acid, or sodium butyrate (NaBu) (95, 96). Dose-response assays were performed to determine the maximum tolerable NaBu dosage for adult *D. simulans*. Doses within a 200-fold range were tested, and the highest dose which did not substantially alter developmental phenotypes was identified (Table S2). This dose was used for all subsequent tests for NaBu impact on CI hatch rates. It is not known what stage(s) of oogenesis are important for conferring Rescue ability upon embryos. Thus, uninfected females were kept on drug food throughout the duration of the experiment except during matings with CI males. Males were incubated and mated on control food only to prevent ingestion of the compound (Figure 2).

**Figure 2.**
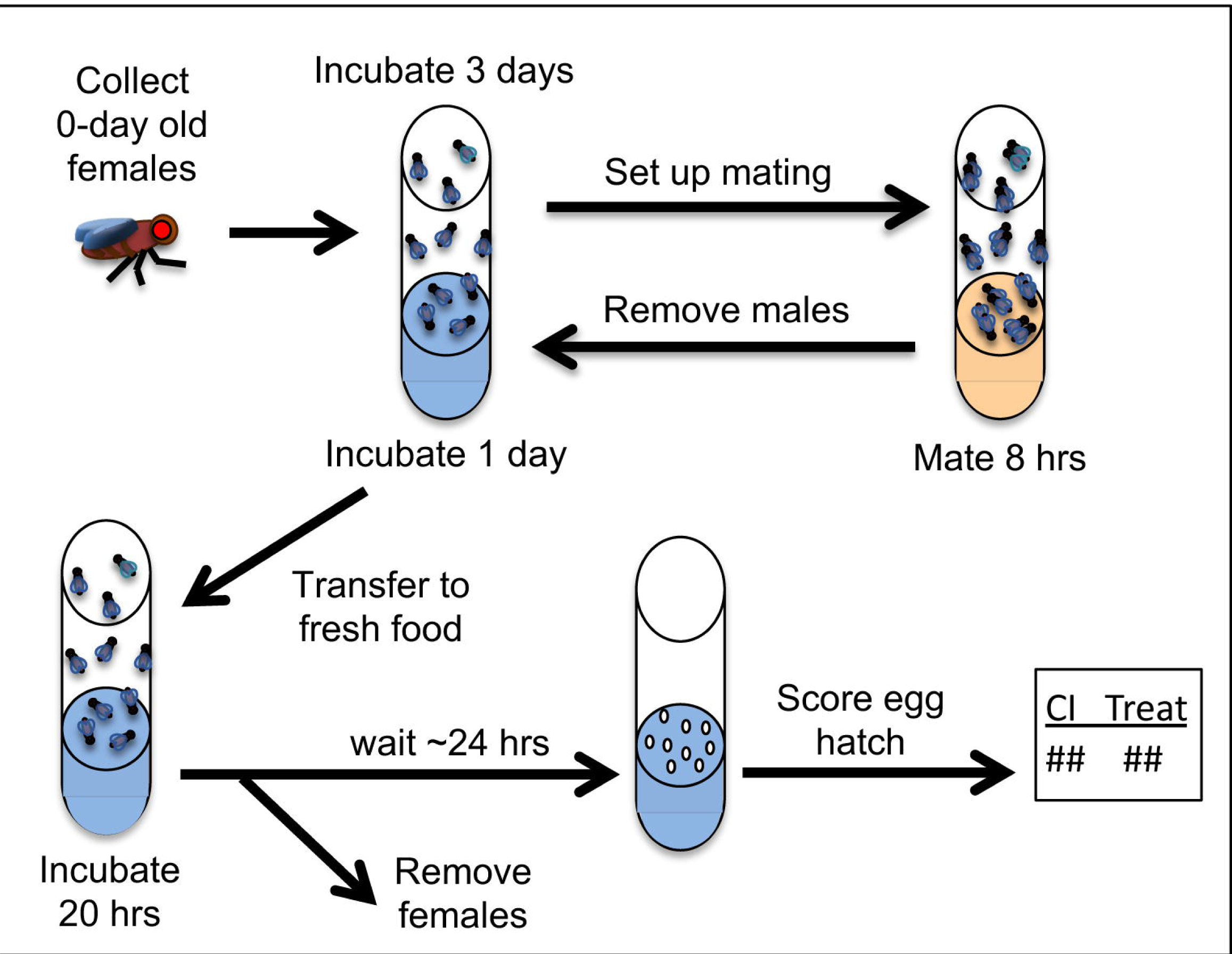
Procedures used for CI- and Rescue-related crosses. Drug feedings were carried out on blue food where specified in the protocol timeline. For details on food preparation, please see Materials and Methods and Additional File 3.

Group mating experiments were performed to determine the impact of NaBu on CI hatch rates associated with *w*Ri and *w*Mel *Wolbachia*. For *w*Ri, the CI hatch rate was 1.6-fold higher for the NaBu treated vials when compared to control food (*p* = 0.0122; adjusted *α* = 0.0125) (Table 2). For *w*Mel, the CI hatch rate on NaBu food was more than twice that of control food conditions (*p* < 0.001 adjusted *α* = 0.0125). To further verify whether the increased egg hatch rates with NaBu feeding were due to unanticipated changes in the mating behavior, single pair matings were also performed. The data confirmed a 94% hatch rate for Rescue crosses, as compared to a 10% hatch rate for CI (*p* < 0.001; adjusted *α* = 0.0083). For the NaBu treatment condition, CI hatch rates were nearly double that of the CI control (*p* < 0.001; adjusted *α* = 0.0083) (Table 2). Taken together, these data indicate that the HDAC inhibitor NaBu induces CI suppression in uninfected females.

**Table 2.**
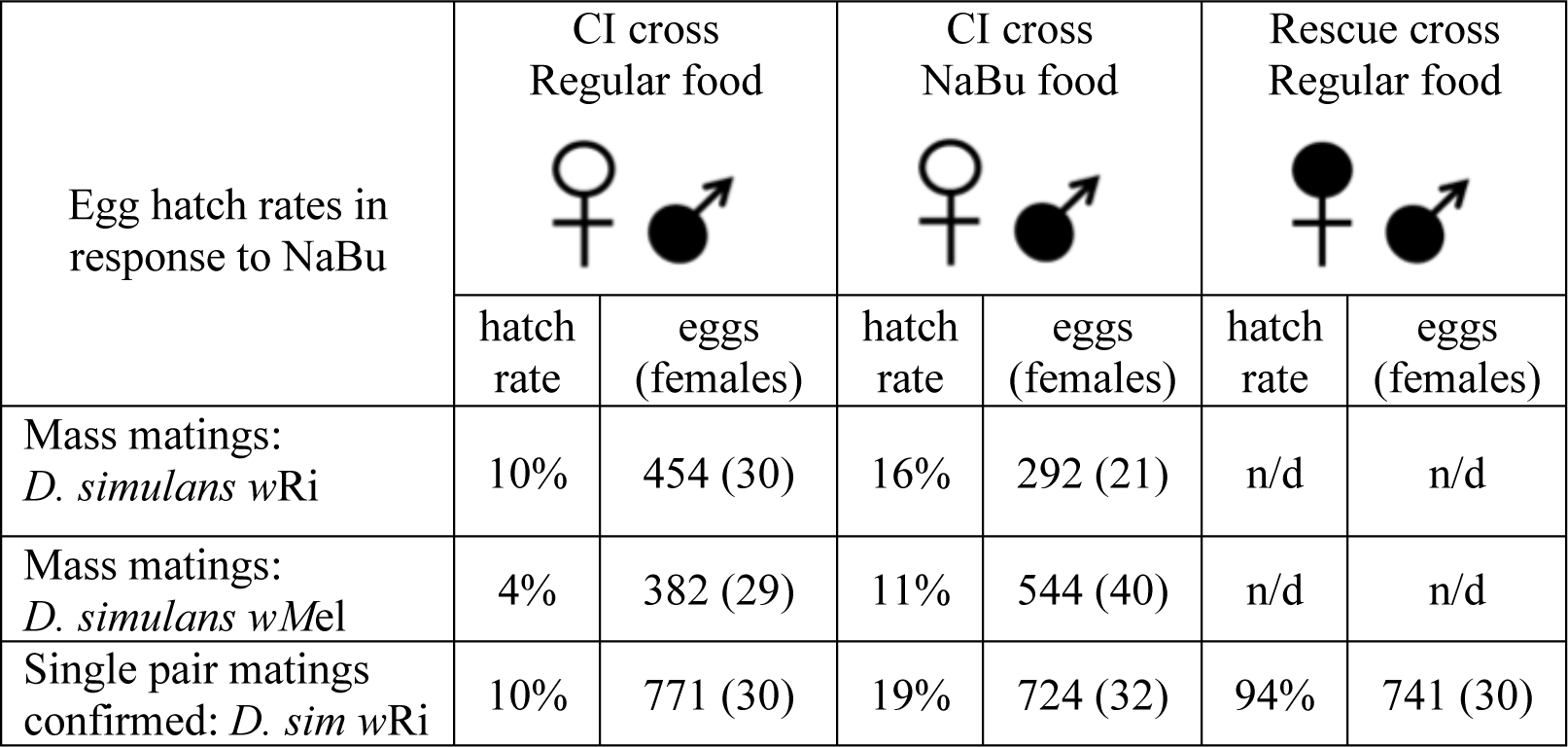
Egg hatch rates from CI-related crosses on NaBu treatment food

| Egg hatch rates in response to NaBu | CI cross<br>Regular food |  | CI cross<br>NaBu food |  | Rescue cross<br>Regular food |  |
| --- | --- | --- | --- | --- | --- | --- |
|  | hatch rate | eggs (females) | hatch rate | eggs (females) | hatch rate | eggs (females) |
| Mass matings:<br>*D. simulans* wRi | 10% | 454 (30) | 16% | 292 (21) | n/d | n/d |
| Mass matings:<br>*D. simulans* wMel | 4% | 382 (29) | 11% | 544 (40) | n/d | n/d |
| Single pair matings confirmed: *D. sim* wRi | 10% | 771 (30) | 19% | 724 (32) | 94% | 741 (30) |

### Scaling up screening of small molecule inhibitors to test for CI suppression

The observation that CI can be partially suppressed by NaBu raises potential questions of whether targeting other host factors could confer CI suppression as well. Assessing this possibility requires screening of more compounds, including sufficient replicates to distinguish CI suppression effects. To this end, we developed a plate-based feeding assay for analyzing small molecule effects on CI hatch rates (76). In the context of 24-well plates, a maximum of 8 wells can be analyzed per condition for CI, CI+treatment and Rescue. Ensuring that results would be relevant to those of a naturally robust CI system, *w*Ri-infected males were used to perform all plate assay matings.

To determine whether chemical feeding in a plate assay format is an effective means of identifying CI suppression, the HDAC inhibitor NaBu was retested across 5 biologically independent plate replicates. All plate replicates indicated consistently higher egg hatch for the CI+NaBu condition (20-30%) than was seen in the CI control condition (11-14%) (*p*-value range: <0.001 to 0.003) (Figure 3A) (Table S3) (Table S4). It is also possible to consider the data from the perspective of plate-based cell screens, where the quality of such assays is typically described in terms of its Z’ factor. Positive Z’ values, ranging between 0 and 1, only result when the average values for the controls are separated by more than 3 times the standard deviation for each treatment (79, 80, 97). According to this analysis, data from the 5 NaBu plate replicates returned Z’ values ranging from 0.76 to 0.89 (Table S5). The CI+NaBu condition also occupied the intermediate “hit” range, consistently distinguishable from the CI control (Figure 3B). This demonstrates that the plate-based feeding assay reproducibly identifies CI-suppressing treatments.

**Figure 3.**
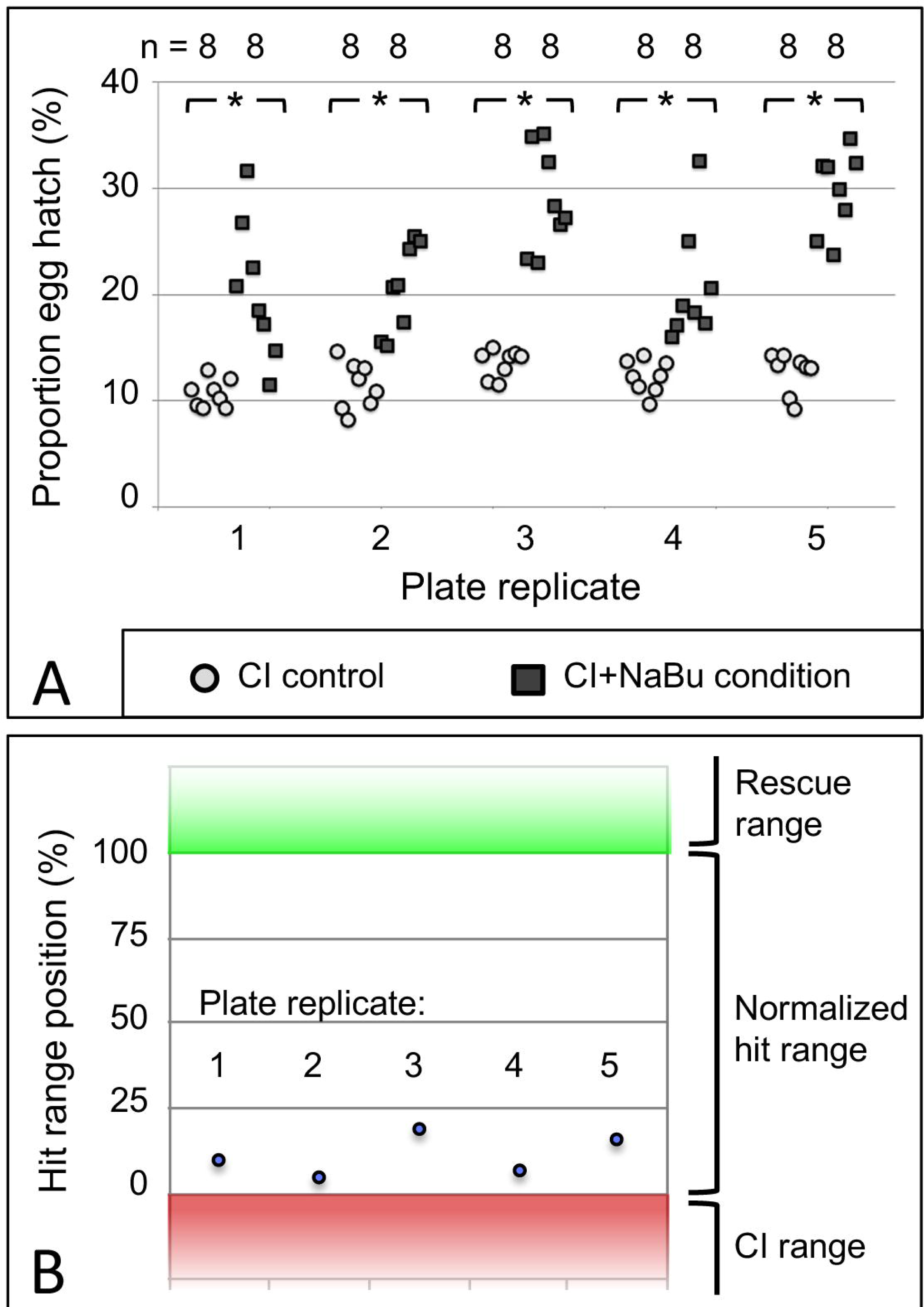
The impact of NaBu treatment on CI egg hatch. (A) Hatch rate data from assay plates using *w*Ri-infected *D. simulans* flies. Each symbol represents data from a single well. * represents *p* < 0.005. (B) NaBu impact on CI, in terms of conventional Z’ analysis. Range boundaries of the CI control (red) and Rescue control (green) are indicated. The normalized “hit range” between controls is shown in white. Blue dots: Average hatch rate for the CI+NaBu condition per screening plate, normalized to the range between CI and Rescue controls. For further details, see Table S4 and Table S5.

To determine the quantity of screening plates required for reproducible identification of a chemical suppressor of CI, the NaBu plate data were statistically analyzed. Data were collated and compared for every crosswise pairing of 5 independent screening plates, using data from 16 wells per condition in each case. Sub-sampling among the 5 plate replicates indicated that data combinations from any 2 plate replicates identified a significant difference between CI and CI+NaBu conditions (*p* < 0.001, n = 10 plate data combinations) (Table S4). To further determine how many wells are required for significance, data were sub-sampled from within each of the paired plate datasets (76). Data from 8 or more wells per condition were sufficient to identify significant differences between CI and CI+NaBu conditions, when setting an alpha value at 0.05 (Figure 4A). Sampling from 11 or more wells per condition yielded a significant difference with alpha set at 0.01 (n = 10 plate data combinations) (Figure 4B). These data indicate that screening 2 chemical assay plates is sufficient to detect chemical suppressors of CI. In addition to corroborating NaBu-induced CI suppression, these plate assay data also created the foundation for testing additional candidate compounds.

**Figure 4.**
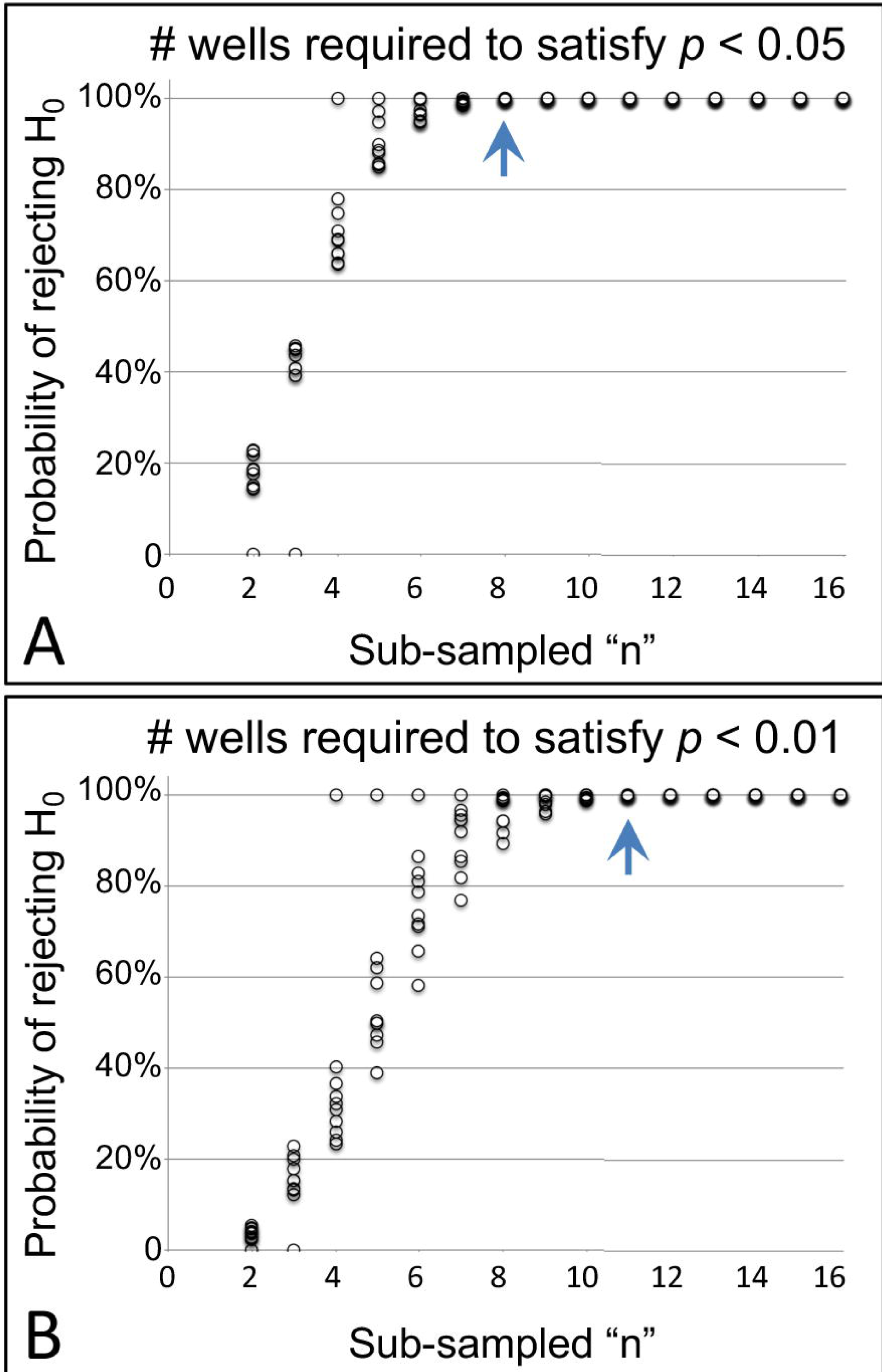
Identifying sufficient sample size in the pate assay format. The graphs show the likelihood of seeing a significant difference in hatch rate between CI and CI+NaBu conditions, as defined by (A) the conventional α-value of 0.05, as well as (B) the more stringent α-value of 0.01. The blue arrow indicates the number of wells at which the probability of rejecting the null hypothesis has reached 99.5% or higher for all sub-sampled datasets analyzed.

### Testing for CI suppression by short-chain fatty acids and protein acetylation modifiers

To further pursue the functional role of NaBu in CI suppression, the basic structure of this compound was considered. Since NaBu is a short-chain fatty acid, this opens the question of whether short-chain fatty acids generally exert CI-suppressing effects. To investigate this possibility, flies were fed with 3 other forms of short-chain fatty acids, specifically acetic, propionic and valeric acid, to test for suppression of *w*Ri-induced CI in *D. simulans.* The doses used for each compound, as well as all others described below, were empirically determined in vial format (Table S6), then tested for impact on CI hatch rates in the context of the plate-based assay. Results from these experiments indicated that acetic acid conferred borderline CI suppression abilities upon uninfected embryos (p = 0.047). Propionic acid and valeric acid had no significant effect on CI hatch rates (Table 3) (Table S7) (Table S8). Thus, CI suppression is not a generalized effect associated with dietary short-chain fatty acids.

**Table 3:**
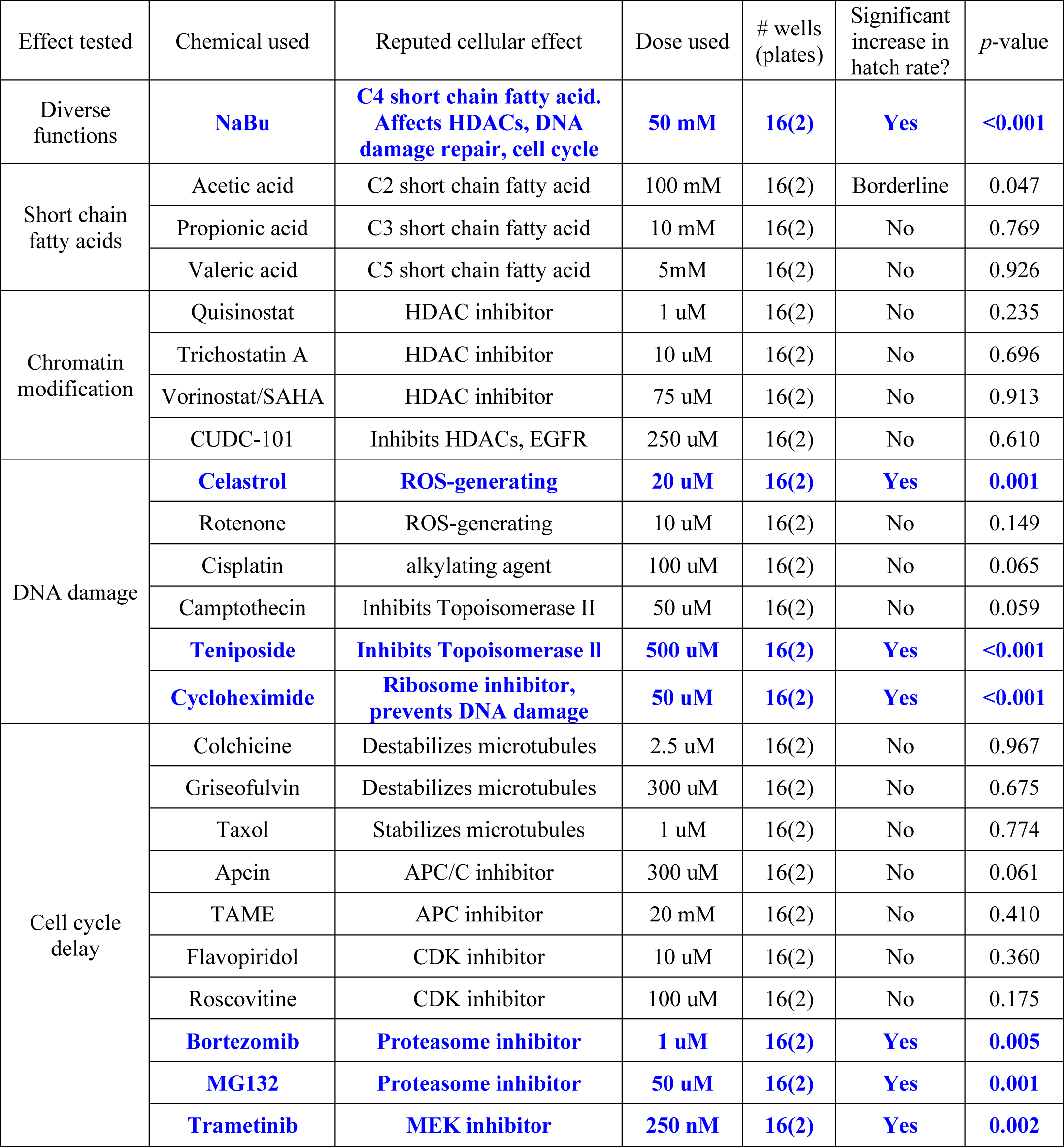
Impact of chemical treatments on CI egg hatch rates for *w*Ri-infected *D. simulans*

| Effect tested | Chemical used | Reputed cellular effect | Dose used | # wells (plates) | Significant increase in hatch rate? | *p*-value |
| --- | --- | --- | --- | --- | --- | --- |
| Diverse functions | **NaBu** | **C4 short chain fatty acid. Affects HDACs, DNA damage repair, cell cycle** | **50 mM** | **16(2)** | **Yes** | **<0.001** |
| Short chain fatty acids | Acetic acid | C2 short chain fatty acid | 100 mM | 16(2) | Borderline | 0.047 |
|  | Propionic acid | C3 short chain fatty acid | 10 mM | 16(2) | No | 0.769 |
|  | Valeric acid | C5 short chain fatty acid | 5mM | 16(2) | No | 0.926 |
| Chromatin modification | Quisinostat | HDAC inhibitor | 1 uM | 16(2) | No | 0.235 |
|  | Trichostatin A | HDAC inhibitor | 10 uM | 16(2) | No | 0.696 |
|  | Vorinostat/SAHA | HDAC inhibitor | 75 uM | 16(2) | No | 0.913 |
|  | CUDC-101 | Inhibits HDACs, EGFR | 250 uM | 16(2) | No | 0.610 |
| DNA damage | **Celastrol** | **ROS-generating** | **20 uM** | **16(2)** | **Yes** | **0.001** |
|  | Rotenone | ROS-generating | 10 uM | 16(2) | No | 0.149 |
|  | Cisplatin | alkylating agent | 100 uM | 16(2) | No | 0.065 |
|  | Camptothecin | Inhibits Topoisomerase II | 50 uM | 16(2) | No | 0.059 |
|  | **Teniposide** | **Inhibits Topoisomerase II** | **500 uM** | **16(2)** | **Yes** | **<0.001** |
|  | **Cycloheximide** | **Ribosome inhibitor, prevents DNA damage** | **50 uM** | **16(2)** | **Yes** | **<0.001** |
| Cell cycle delay | Colchicine | Destabilizes microtubules | 2.5 uM | 16(2) | No | 0.967 |
|  | Griseofulvin | Destabilizes microtubules | 300 uM | 16(2) | No | 0.675 |
|  | Taxol | Stabilizes microtubules | 1 uM | 16(2) | No | 0.774 |
|  | Apcin | APC/C inhibitor | 300 uM | 16(2) | No | 0.061 |
|  | TAME | APC inhibitor | 20 mM | 16(2) | No | 0.410 |
|  | Flavopiridol | CDK inhibitor | 10 uM | 16(2) | No | 0.360 |
|  | Roscovitine | CDK inhibitor | 100 uM | 16(2) | No | 0.175 |
|  | **Bortezomib** | **Proteasome inhibitor** | **1 uM** | **16(2)** | **Yes** | **0.005** |
|  | **MG132** | **Proteasome inhibitor** | **50 uM** | **16(2)** | **Yes** | **0.001** |
|  | **Trametinib** | **MEK inhibitor** | **250 nM** | **16(2)** | **Yes** | **0.002** |

NaBu is best known for its impact on chromatin structure and has been credited with suppressing HDACs 1-5 and 7-9, representing the entirety of class I and IIa HDAC enzymes (93, 94). To determine whether maternal HDAC function affects *w*Ri-induced CI, an array of HDAC inhibitors was dose-optimized (Table S6) and tested in the plate-based assay. The selected inhibitors included the class I and II HDAC inhibitors quisinostat (98) and trichostatin A, as well as the pan HDAC inhibitor vorinostat (SAHA) (93, 94). The inhibitor CUDC-101, which targets class I and II HDAC as well as growth factor receptors, was also tested (99). Remarkably, none of these HDAC inhibitors exerted significant impact on CI hatch rates (Table 3) (Table S7) (Table S8). To confirm that the fly stocks and assay parameters were still performing as expected, the effect of NaBu was retested in the plate-based format. The data confirmed that CI suppression by NaBu was still significant, with a *p*-value below 0.001 (Table 3) (Table S7) (Table S8). Overall, these results do not support HDAC inhibition and associated chromatin remodeling as a generalized mechanism for CI suppression.

### Testing modifiers of DNA damage for maternal CI suppression effects

NaBu has also been shown to promote DNA repair, in part by indirectly increasing acetylation of histone H4 (100-102). To test whether maternal DNA repair processes affect *w*Ri-induced CI, an array of inhibitors was pursued. To activate the DNA repair response, agents that induce oxidative DNA damage were selected, specifically celastrol and rotenone (103-105). The alkylating agent cisplatin, which also generates reactive oxygen, was included as well (106-108). Topoisomerase inhibitors were also used, including camptothecin, which prevents re-sealing of single stranded nicks by topoisomerase I, and teniposide, which prevents removal of topoisomerase II from DNA and induces degradation of the enzyme (109-111). As the ribosome inhibitor cycloheximide has been reported to prevent formation of single- and double-stranded DNA breaks (112, 113), this drug was also tested.

Using optimized doses (Table S6), the plate-based feeding assay indicated CI suppression for half of the treatment conditions used. A significant increase in hatch rate was observed for uninfected females exposed to celastrol, teniposide and cycloheximide, all associated with *p*-values of 0.001 or less (Table 3) (Table S7) (Table S8). These data open a possible role for maternal processes that prevent and repair DNA damage in conferring CI suppression upon uninfected embryos.

### Testing the impact of cell-cycle timing on maternal CI suppression

Exposure to NaBu has been shown to slow cell cycle timing (114, 115). To test the effect of maternal cell cycle timing on *w*Ri-induced CI, uninfected *D. simulans* were exposed to an array of complementary inhibitors. In attempt to slow the progression of mitosis by altering microtubule dynamics, the microtubule destabilizers colchicine and griseofulvin, as well as the microtubule stabilizer taxol were tested (116, 117). To slow anaphase onset and exit from mitosis, inhibitors of the anaphase promoting complex, apcin and TAME, were used (118, 119). To inhibit the progress of mitosis and the cell cycle overall, the Cyclin dependent kinase inhibitors flavopiridol and roscovitine were used(120-122), as well as the proteasome inhibitors bortezomib and MG132 (123, 124). To stall general re-entry into the cell cycle, the MAPKK (MEK) inhibitor trametinib was also used (125, 126).

After identifying appropriate doses (Table S6), chemical manipulators of cell cycle timing were tested for CI suppression. The plate assay data indicated significantly increased CI hatch rates for bortezomib, MG132, and trametinib-fed females compared to control (*p*-value range: 0.001-0.005) (Table 3) (Table S7) (Table S8). As cell cycle delays are a recognized consequence of DNA damage, resulting from checkpoint activation that allows damage repair (127), these data are consistent with a possible role for altered embryonic cell cycle timing in suppression of CI.

### Re-testing CI-suppressing compounds against transinfected *D. simulans*

If the compounds that suppress *w*Ri-induced CI act upon a network of conserved, Rescue-related maternal interactions, the effects would be expected to be applicable to other host-strain combinations. To this end, the chemicals identified above as hits were analyzed for suppression of *w*Mel-induced CI as well, using flies from a transinfected stock population (73). Specifically, NaBu, celastrol, cycloheximide, teniposide, bortezomib, MG-132, trametinib, and the initially borderline hit acetic acid (Table 3) were retested in the plate assay format. The same dosing and procedures were used as above, with the only difference being that *w*Mel-infected males were used for CI induction.

The results indicated that NaBu, celastrol, and cycloheximide significantly elevated CI hatch rates for *w*Mel-induced CI (*p*-value range: 0.011-0.013) (Table 4) (Table S9) (Table S10). By contrast, teniposide, bortezomib, and MG-132 treatments exhibited borderline CI suppression effects (*p*-value range 0.041-0.047). Trametinib and acetic acid did not induce any significant effects (Table 4) (Table S9) (Table S10). This outcome distinguishes cellular responses associated with certain DNA damage and/or cell cycle timing regulators as general contributors to maternal suppression of CI in *D. simulans.*

**Table 4:** Impact of chemical treatments on CI egg hatch rates for *w*Mel-infected *D. simulans*

| Effect tested | Chemical used | Reputed cellular effect | Dose used | # wells (plates) | Sig increase in hatch rate? | *p*-value |
| --- | --- | --- | --- | --- | --- | --- |
| Diverse functions | **NaBu** | **C4 short chain fatty acid. Affects HDACs, DNA damage repair, cell cycle** | **50 mM** | **16(2)** | **Yes** | **0.011** |
| Short chain fatty acids | Acetic acid | C2 short chain fatty acid | 100 mM | 16(2) | No | 0.408 |
| DNA damage | **Celastrol** | **ROS-generating** | **20 uM** | **16(2)** | **Yes** | **0.013** |
|  | **Cycloheximide** | **Ribosome inhibitor, prevents DNA damage** | **50 uM** | **16(2)** | **Yes** | **0.013** |
|  | Teniposide | Inhibits Topoisomerase II | 500 uM | 16(2) | Borderline | 0.041 |
| Cell cycle delay | Bortezomib | Proteasome inhibitor | 1 uM | 16(2) | Borderline | 0.047 |
|  | MG132 | Proteasome inhibitor | 50 uM | 16(2) | Borderline | 0.047 |
|  | Trametinib | MEK inhibitor | 250 nM | 16(2) | No | 0.096 |

### Testing combined pathway effects for suppression of CI

To further test the extent to which suppression of CI under these treatments is due to a shared network of pathway functions, a dual chemical treatment strategy was pursued. We focused on hits that elicited the most robust CI suppression across both systems tested, namely: NaBu, celastrol, and cycloheximide. Additional treatment combinations included compounds that exerted more modest effects, namely: teniposide, bortezomib, and MG-132. As for the single drug trials, dose response curves were carried out for all pairwise chemical treatments on uninfected *D. simulans* flies (Table S11). Nearly all treatment combinations required reduced drug dosing, as compared to singly administered treatments (Table S6). The cycloheximide/bortezomib combination was the only case in which original treatment doses for both compounds could be tolerated additively, without adverse effects on fecundity, egg hatch or larval development (Table S11).

After uninfected *D. simulans* females were treated with dual drug combinations, their egg hatch rates were compared against females raised on control food in the plate assay format. To ensure that the results would be representative of natural CI, *w*Ri-infected males were used to induce CI in this series of experiments. The results indicated that the cycloheximide/bortezomib combination significantly increased the CI hatch rate as compared to the CI control (*p* = 0.006) (Table 5) (Table S12) (Table S13). No other paired chemical treatments induced CI suppression (Table 5). It is possible that loss of CI suppression effects is attributable to reduced combinatorial doses of otherwise effective compounds. Regardless, the data indicate CI suppression by compounds that are traditionally associated with manipulation of protein synthesis and protein turnover.

**Table 5.** Impact of chemical combinations on CI egg hatch rates for *w*Ri-infected *D. simulans*

| Effect tested | Chemical used | Dose used | # wells (plates) | Significant increase in hatch rate? | *p*-value |
| --- | --- | --- | --- | --- | --- |
| Diverse functions/<br>DNA damage | Nabu/Celastrol | 25 mM / 10 µM | 16(2) | No | 0.119 |
| Diverse functions/<br>DNA damage | Nabu/Cycloheximide | 25 mM / 25 µM | 16(2) | No | 0.817 |
| Diverse functions/<br>Cell cycle delay | Nabu/MG132 | 25 mM / 25 µM | 16(2) | No | 0.201 |
| DNA damage | Teniposide/Celastrol | 250 µM / 500 µM | 16(2) | No | 0.287 |
| DNA damage | Celastrol/Cycloheximide | 10 µM / 25 µM | 16(2) | No | 0.565 |
| **DNA damage/**<br>**Cell cycle delay** | **Cycloheximide/Bortezomib** | 50 µM / 1 µM | **16(2)** | **Yes** | **0.006** |
| DNA damage/<br>Cell cycle delay | Teniposide/MG132 | 250 µM / 25 µM | 16(2) | No | 0.264 |

## DISCUSSION

This study was designed to inform mechanisms associated with Rescue in robust CI systems, endogenous to non-model organisms. A strong case is currently being made for involvement of *Wolbachia* Cif proteins in induction of CI and Rescue, based on analysis of *D. melanogaster* and yeast models (60-62, 65, 66). This study, using natural and transinfected *D. simulans*, add to the complex biological underpinnings of embryonic lethality by indicating that maternal contributions to Rescue are also possible. Demonstrating CI suppression through chemical feeding of uninfected females, without invoking any effect or contribution by *Wolbachia*-supplied antitoxins, opens many questions regarding how Rescue functions across *Wolbachia-*host systems. Fundamental to these questions is whether a core set of maternal mechanisms in insects can act to suppress CI across systems. Fortunately, the experimental framework presented here enables broad investigation of diverse CI-Rescue systems in the future, including that of non-*Wolbachia* endosymbionts like *Cardinium* (128-130).

Despite the reproducibility and statistical significance of CI-suppression effects by multiple drugs and drug combinations, the limitations of the current work are reflected by CI hatch rates which did not exceed 3 times that of CI-control hatch rates. The disparity between the 90%+ egg hatch frequencies of *Wolbachia-*induced Rescue and that of chemical CI suppression can be due to a variety of factors or experimental limitations. One important aspect could be the absence of *Wolbachia-*supplied Cif proteins. It is possible that maternal mechanisms act as a supplement to, in parallel with, or in coordination with Cif functions to a substantial extent. There are no known chemical treatments that will mimic the full array of predicted functions associated with Cif proteins at this time (62, 63). Our attempt to alter broader functional networks of maternal proteins by use of multiple inhibitors was met with limited success, possibly due to dosage considerations, limited by systemic tolerances for dual treatments.

When interpreting the CI suppression data yielded by the current chemical screen, the technical limitations inherent to the method itself are important to consider. Whole body feedings lack the time and tissue-specific nuance afforded to *Wolbachia in vivo.* While dosing within the food was standardized for this study, it is not possible to control for local dosing to tissues/cells of the recipient organism. This is due to differences in ingestion, absorption, efflux, metabolism, and/or excretion rates, which are expected to vary in association with each cell type and each drug. A feeding assay also creates the possibility for side effects due to host microbiome impacts. Initial attempts to run this screen using standardized micro-injections were curtailed by observations that injected *D. simulans* flies stop laying eggs. Working with flies under anexic or gnotobiotic conditions presents its own set of complications (131). For these reasons, seeing a response from this chemical screen is informative, whereas the lack of a response is not. Distinct from most drug screens, whose interpretations are limited by typically detrimental health effects on the test subject, the output of this screen leads to increased survival of otherwise ill-fated embryos. Demonstration of CI suppression adds a role for maternally contributed factors to otherwise antitoxin-attributed Rescue effects.

This study was designed to address potential contributions of chromatin remodeling, DNA damage repair and cell cycle timing to the mechanism of Rescue. Though little support was evident for chromatin remodeling in CI suppression, impacts on DNA integrity and cell cycle timing conferred significant increases in CI egg hatch. As such, our results at present do not readily distinguish between existing models of CI and Rescue, but instead opens consideration of networked models that may better reflect the cell biology of CI. DNA damage and cell cycle timing are well known to be intrinsically connected, since DNA damage triggers a checkpoint mechanism that arrests the cell cycle (127, 132). Variation upon functions of the ubiquitin-proteasome system may further link these processes. DNA damage repair is facilitated by ubiquitination of histones and DNA repair pathway proteins, followed by their deubiquitination upon completion of the repair (133-135). Ubiquitination and degradation of cyclins and other regulators is also fundamentally required for cell cycle progression (132, 136). Expression studies in CI-inducing *Cardinium* have implicated the ubiquitin-proteasome system, as well as DNA repair processes (128). The CI-suppressing compounds identified in this study accordingly reflect this continuum of function.

The use of well-known compounds in this study provides access to vast literature for interpreting reproducible CI suppression effects. One question raised by this work is what distinguished the compounds that affected egg hatch for both *w*Ri and *w*Mel-induced CI, from those that affected *w*Ri-induced CI only (Figure 5). Studies in mammalian systems and cell lines indicate that these compounds consistently induce cell cycle arrest, but exert variable impacts on protein turnover, depending upon the compound (Additional File 3). Neither of those functional profiles, as reported in the literature, currently aligns with the strain-specific differences observed in CI egg hatch outcomes (Figure 5).

**Figure 5.**
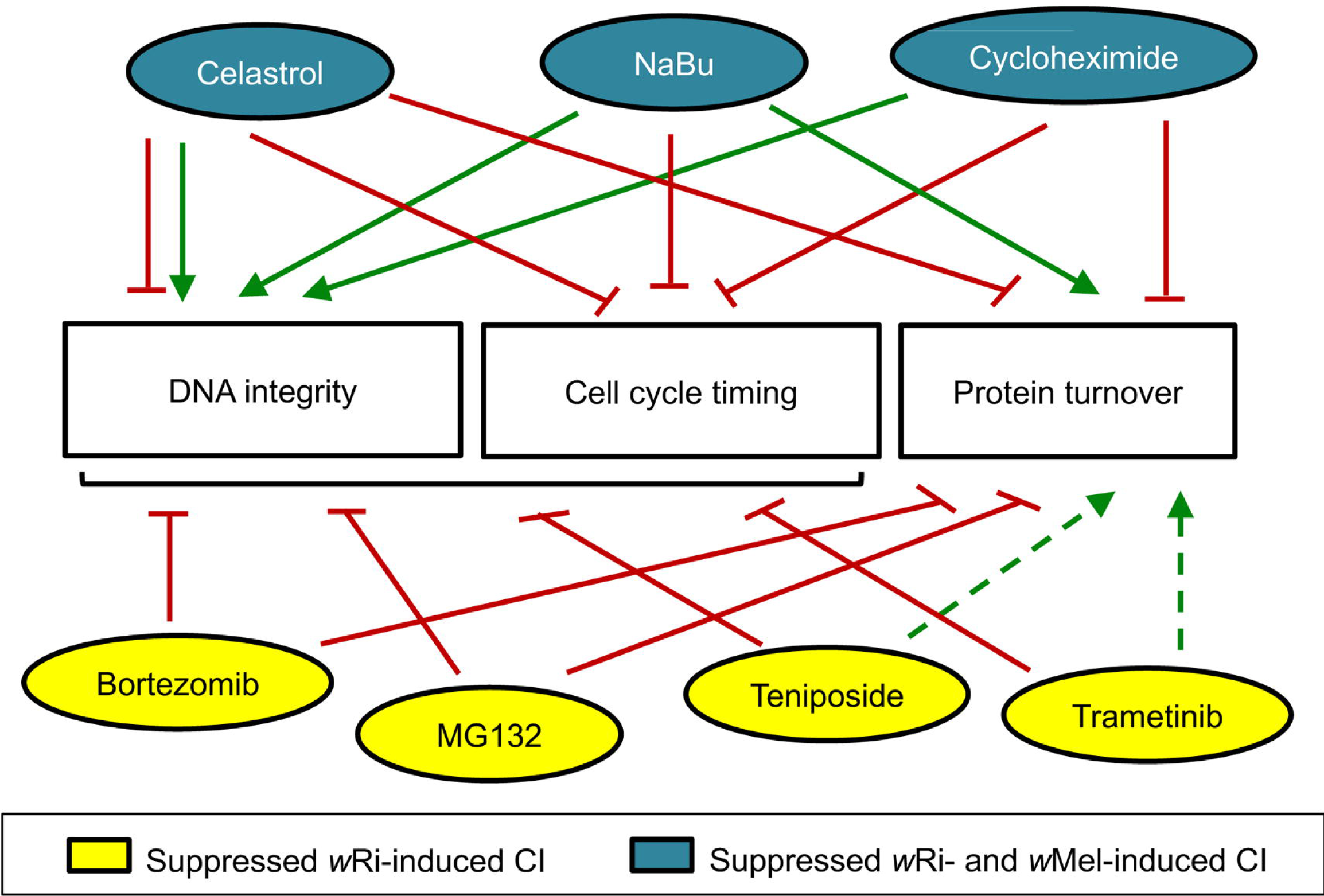
Summary of maternal impacts that significantly increased *D. simulans* CI hatch rates. This information is based upon available literature, summarized in Additional File 3. Green arrows: positive impact. Red lines: negative effect. Dotted lines: interpretation based upon partial datasets. Bracket: includes multiple categories.

Clearer associations are evident between existing literature and CI suppression outcomes from this study from the perspective of DNA integrity. The 4 compounds that suppress *w*Ri-induced CI are known inducers of DNA damage, whereas the 3 compounds that suppress both *w*Ri- and *w*Mel-induced CI, reportedly support DNA integrity (Figure 5) (Additional File 3). Though celastrol can exert detrimental impacts on DNA (137, 138), it has also been shown to suppress radiation-induced damage (103, 104). Cycloheximide treatments prevent formation of single- and double-strand DNA breaks (112, 113). Lastly, NaBu protects DNA integrity by up-regulating antioxidant pathways and by facilitating DNA repair (100, 101, 139). This suggests that DNA integrity is a dynamic, focal aspect of CI suppression by different *Wolbachia* strains in *D. simulans*. The *Wolbachia*-encoded, toxin-protein CidB was recently linked to direct inhibition of karyopherin function, involved in eukaryotic cytoplasm-to-nucleoplasm transport, with the implication that CidB may prevent access of chromatin interacting factors to paternal chromatin (66). Future analyses of CI and Rescue, from the perspective of both host and microbe promise to be informative in elucidating the molecular basis of this ecologically relevant mechanism.

## AUTHOR CONTRIBUTIONS

AJMZM, ADAS, JGA, SAS, AF, JH, SC and LRS designed the experiments.

AJMZM, ADAS, SAS, SC and LRS supervised the experiments.

AJMZM, ADAS, JGA, SAS, AF, JH and SC conducted the experiments.

AJMZM, ADAS, JH, SC and LRS wrote the manuscript.

AJMZM, ADAS, JGA, SAS, AF, JH, SC and LRS reviewed the manuscript.

## ACKNOWLEDGEMENTS

We thank the Serbus lab, past and present, William Sullivan, Michael Turelli, the FIU Department of Biological Sciences, and the FIU Biomolecular Sciences Institute for helpful discussions and support.

## DATA AVAILABILITY STATEMENT

The datasets generated by this study can be found in the accompanying Supplementary Tables.

