## Additional File 1 for "A Role for Maternal Factors in Suppressing Cytoplasmic Incompatibility"

**Protocol: Plate-based chemical feeding assay**

**Preparing chemicals:**

*POWDERS*

Plan to just mix/add this in on the day of the experiment.

Sodium Butyrate (NaBu): Sigma cat# B5887-1G

Mix 110mg powder directly into 20mL of melted (then cooled down) fly food

Final drug concentration in food: 50mM

*FREEZER STOCKS (TYPICAL)*

Prepare stock solutions in advance and refrigerate/freeze down in 1mL aliquots.

These can be used (freeze/thawed) up to 4-5 times.*

Bortezomib: Adipogen cat# PS-341 (we ordered this thru Fisher)

Note this drug is light-sensitive. Must be stored in the dark. Upon use, keep vials/plates wrapped in foil

Stock: 1mM suspended in DMSO

Mixture in Food (20 mL): add 20μL stock solution

Final drug concentration in food: 1μM

Celastrol: Sigma cat# C0869-10MG

Stock: 5mM suspended in DMSO

Mixture in Food (20 mL): add 80μL stock solution

Final drug concentration in food: 20μM

Cycloheximide: Sigma cat# C7698-1G

Stock: 20mM suspended in DMSO

Mixture in Food (20 mL): add 100μL stock solution

Final drug concentration in food: 50μM

MG-132: Sigma cat# SML1135-5MG

Stock: 50mM suspended in DMSO

Mixture in Food (20 mL): add 20μL stock solution

Final drug concentration in food: 50μM

Teniposide: Sigma cat# SML0609-50MG

Stock: 100mM suspended in DMSO

Mixture in Food (20mL): add 100μL stock solution

Final drug concentration in food: 500μM

*FREEZER STOCKS (NON-TYPICAL)*

Trametinib: Fisher cat# NC0592237

This compound is highly sensitive to freeze/thaw cycles.

Stock: Arrives as a 10mM stock in DMSO

Plan to create 10+ aliquots that each contain 10μL, ok to re-freeze at -20C.

When removing a tube for use in an experiment, do not re-freeze/re-use again though.

On day of use: Dilute 5μL of the 10mM stock into 995μL of DMSO to make a 50μM working stock

Mixture in Food (20mL): add 100μL working stock solution

Final drug concentration in food: 250nM

**Preparing plastics:**

Get out three 24-well plates (Corning cat# 3738)

Two of these plates will be physically unaltered, and used for dispensing food (left plate image)

The third plate has to be altered, with the bottom part converted into a “Perfusion-friendly lid”

Get out a crescent wrench, and tighten it around a small area of the short side wall

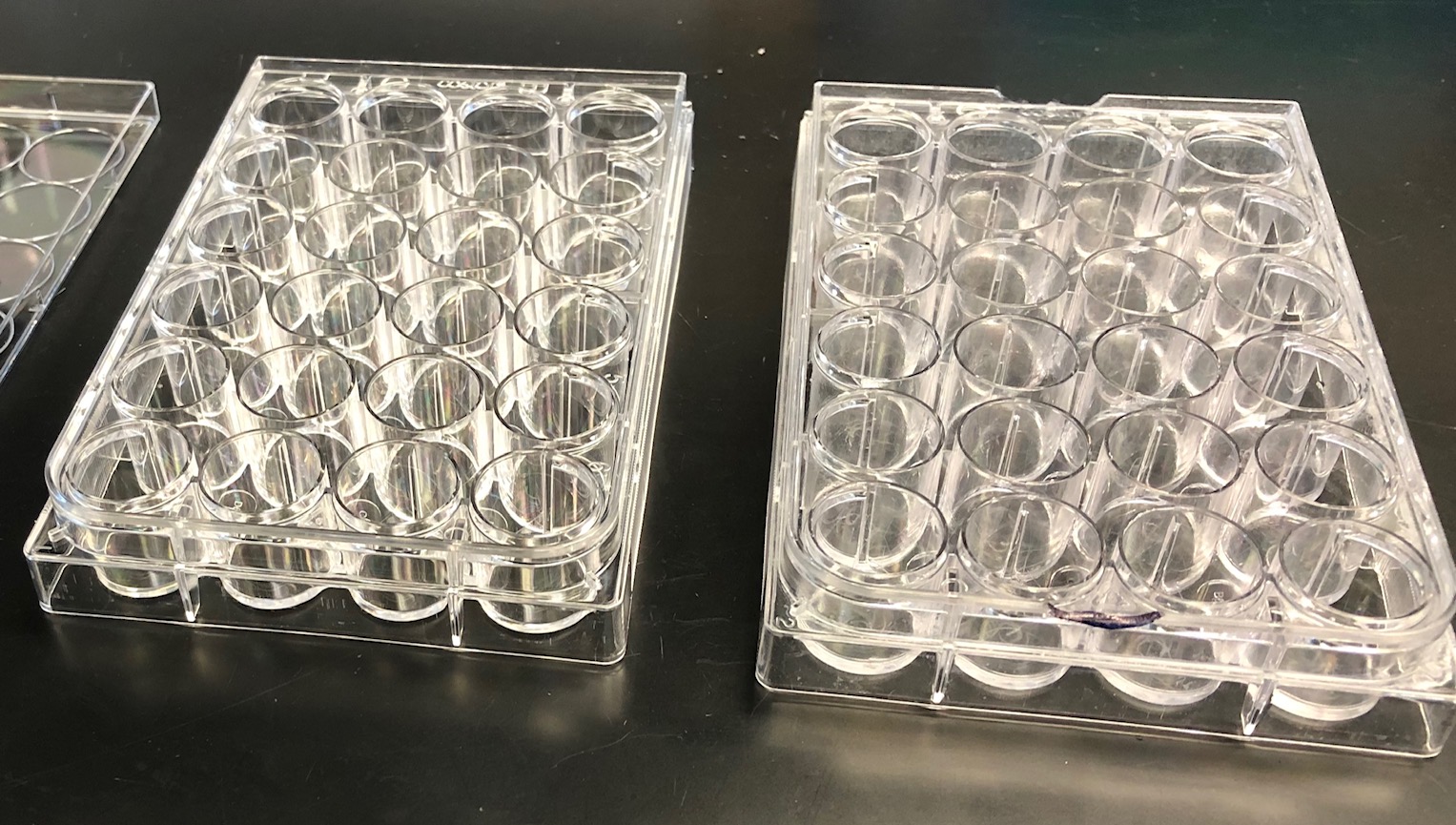

Standard plate	Perfusion-friendly lid

Flex the crescent wrench outward to snap off a tiny bit of the short side wall

Repeat on the second short side, so that both sides of plate will have little holes. (right side plate image)

Creating little holes on each short side of the lid will allow CO_2_ to flow thru there later.

Also cut off the tips from at least four 1mL pipet tips, leave in the fume hood.

Leave a 1000uL pipetting device in the hood.

**Preparing plate content:**

Weigh out 6.4mg Brilliant Blue G powder (Acros Organics cat# 191480250), set aside.

Also get out a 100mL clean beaker and two 60mL beakers.

Also get out an aliquot of the chemical stock to be used from storage (fridge or freezer).

If the chemical is frozen, let it sit out at room temp in hood during fly food prep (below)

Retrieve prepared Drosophila rearing media from the 4°C fridge.

Spoon out 60mL (approx) into a 100mL beaker (ok to smash food down a bit while adding)

Put food beaker in the microwave, and heat it. **Wait there and watch, to prevent food volcano.*

As soon as you see the food start to bubble:

Take beaker out of microwave, stir, put back in microwave.

Repeat cycles of heating and stirring until all food is melted and uniform in consistency.

Using our microwave, this = about 1 min of cook time, interrupted 2-3x with stirring.

While food is still very warm and runny, add 6.5mg Brilliant Blue G powder directly to food

Stir dye into the food using a spoon-shaped spatula (Fisher cat# 03-990-240)

Make sure there are no streaks in the food while stirring.

If streaks cannot be stirred out, microwave 10-15 sec more until bubbly, resume stirring.

Split the 60mL of blue food into two new (60mL) beakers—put 20mL into one, 40mL into other.

Note: by this time the food will still be runny, but more moderate in temperature.

Put the 20mL blue food into the fume hood. This will be the **“treatment”** food.

Pipet (thawed) chemical stock solution onto the surface of the fly food.

Stir the chemical stock into the food, VERY thoroughly, for at least 30 seconds.

This means stir, then scraping down the sides, and scrape across the bottom,

Then stir more, etc. Important to plan on repeating a couple cycles of this.

Then immediately pipet the food into the appropriate wells, as described below.

If the food thickens while dispensing, re-melt again, only very briefly, in the microwave.

Put the 40mL blue food into the fume hood. This will be the **“control”** food.

Pipet appropriate amount of water or DMSO to equivalent dilution used for treatment food above.

When preparing this, use 2X the volume that was used for the chemical.

i.e. if 20uL Bortezomib stock was added to treatment beaker,

then you would plan to add 40uL of DMSO to the control food beaker.

Stir the water or control DMSO into the food, VERY thoroughly, for at least 30 seconds.

This means stir, then scraping down the sides, and scrape across the bottom,

Then stir more, etc. Important to plan on repeating a couple cycles of this.

Then immediately pipet the food into the appropriate wells, as described below.

**Dispensing content into plates:**

Make sure to dispense fly food into the center of the well. (Can’t count egg lay on side of well.)

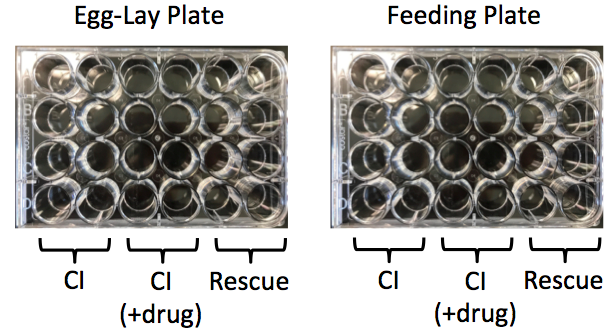
Plate format will run as follows:

Feeding Plate	Egg-Lay Plate

Using a 1mL pipette tip w/ tip cut open to accommodate fly food, deliver **~700-800 uL** per well

-apply “treatment” food to 2 center columns (8 wells per plate x2 plates)

-apply “control” food to the outer 2 columns on the left and right sides of each plate

(16 wells per plate x2 plates)

If the food starts accumulating within the pipet tip to the point that it approaches the pipet barrel

Eject the pipet tip, then continue dispensing food with a new pipet tip.

Cool/air dry in the fume hood, covered lightly with cheesecloth, for 2 hrs

This wait time will prevent condensation that otherwise happens later in the plates.

-At end of this process, you have two plates set up.

Identify which of the two plates came out the smoothest, with no fly food stuck on the walls.

Label this plate “Egg lay Plate”

Place the plate cover back on the EGG LAY PLATE.

Store this plate in a zip lock bag, and store the sealed bag at 4°C .

The egg laying plate will be used on DAY 4.

The other, remaining plate should be labeled FEEDING PLATE.

This will be used starting today, DAY 0.

**Feeding/mating schedule:**

This is assuming the incubator is set for lights out at 10pm, lights back on at 1pm next day.

*Flies are reared as per the separate protocol below.

**DAY 0-- Begin plate set-up (must be completed by early evening)**

Take boxes of bottles to be used out of the incubator between 9-9:30am.

*Ok to leave boxes out on the benchtop from now until fly collection is completed.*

Clear ALL existing flies out of the bottles.

(After tray cleared, recheck all bottles to be sure and eliminate any/all stragglers)

Within the next 8 hrs (ideally 6 hrs or less), collect all newly eclosed flies.

We typically collect at 2pm, and collect again at 5pm if not enough flies were available.

Note that fly collection at multiple timepoints is important if more flies are needed

Cannot let bottles sit/accumulate flies 8+ hours, that = lots of non-virgin females.

Collection goals for running a single plate assay:

Need a total of 160 uninfected females (plan to add these to the plate today)

Need 80 *w*Ri-infected females (plan to add these to plate today)

Need 240 *w*Ri*-*infected males (plan to save these for later!!)

Store the males in 4 vials containing regular food, with maximum 60 males/vial.

Bring chemical feeding plate and the perfusion-friendly plate lid over to the fly pushing area.

Set aside chemical feeding plate for a few minutes—don’t put any sleeping flies in here.

Plan to load flies, pre-separated into piles of 10 flies/well, into the perfusion-friendly plate lid.

**critical to prevent flies from getting stuck in the food while asleep.*

Check plate lid orientation to ensure fly type and food condition will correctly match up later.

Use a funnel to quickly dispense 10 females to the middle of each well area on plate lid.

Add uninfected females to left 4 columns (16 wells, for CI and CI+treat wells)

Add *w*Ri-infected flies to the right 2 columns (8 wells, for Rescue wells)

Have a second gas pad handy to apply CO_2_ to surface of wells if flies wake while doing this

If flies kick/agitate, invert gas pad, touch to top of plate lid to re-anesthetize.

Carefully place the chemical feeding plate on top of the perfusion-friendly lid to close the plate.

Put bench tape along 2 long sides of plate to secure perfusion-friendly lid onto food section.

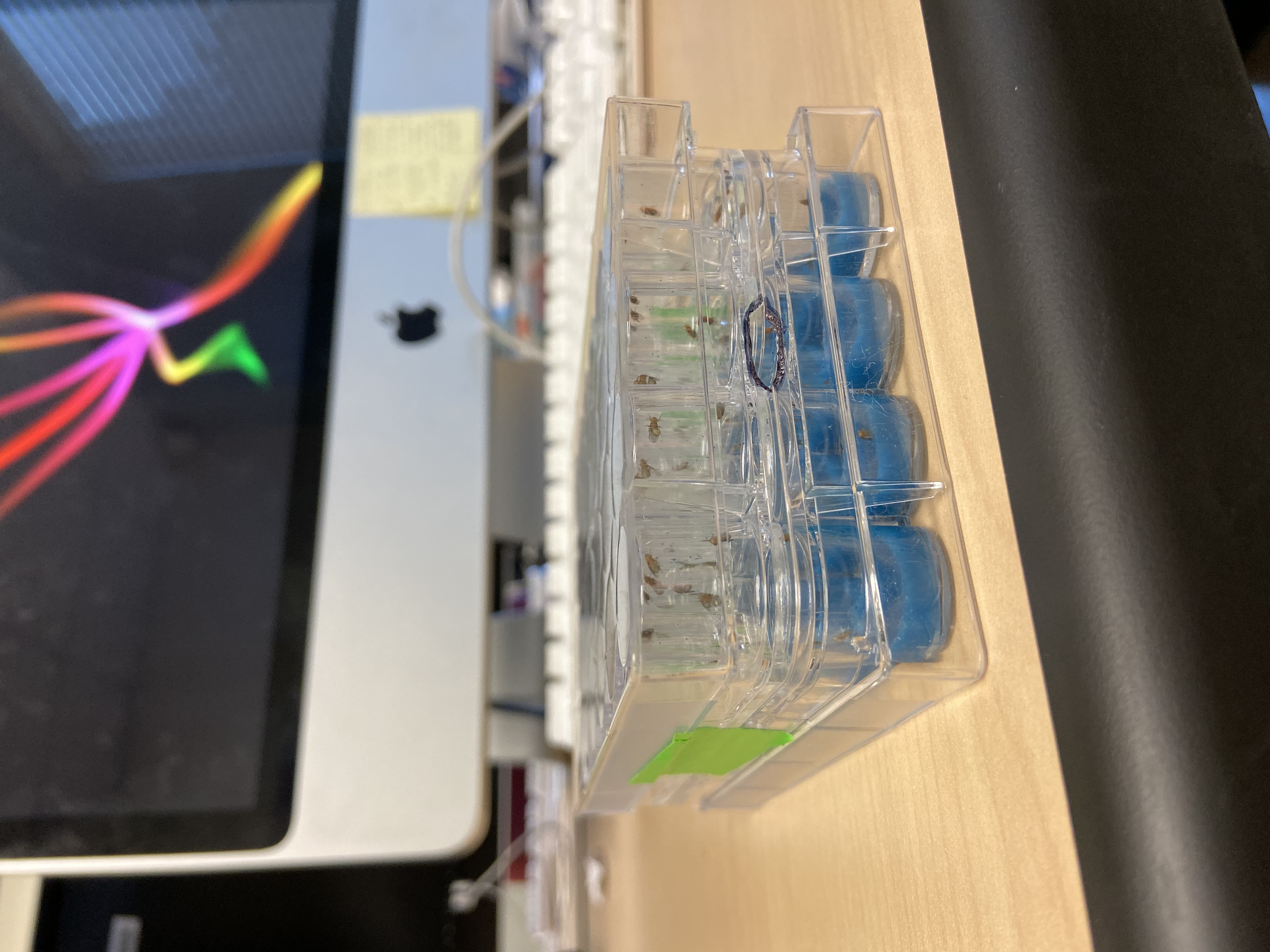
**Make sure that alignment of the perfusion-friendly lid and the feeding plate is precise,*

*in both directions, so flies can’t crawl from one well to another.*

Wait until all the flies are fully awake before flipping plate over (*prevents flies getting stuck*)

Place the plate in the 25°C fly incubator

Use a middle shelf for the best moisture control.

AVOID bottom shelf of the incubator.

That seems to drive condensation.

Let the plate incubate at 25°C for 3 days.

**DAY 2 Create a new plate food plate (to use for mating step tomorrow)**

Prepare a 24-well plate with standard lab fly food.

Melt 40mL food in the microwave, stirring as you go.

It’s ok to just use regular/untreated food, as blue dye is not needed for this.

Using a 1mL pipette tip w/ tip cut open to better pick up fly food, deliver ~700-800 uL to each well

If the food starts accumulating within the pipet tip to the point that it approaches the pipet barrel

Eject the pipet tip, then continue dispensing food with a new pipet tip.

Cool/air dry in the fume hood, covered lightly with cheesecloth, for 2 hrs

Put the plate in a ziplok bag, then store the plate in the 4°C fridge.

The plate you just made is hereafter referred to as the “mating plate.”

Using this tomorrow will ensure that CI males never see any drug, even during the mating step.

**DAY 3 Set up the mating** (**anytime between 10am-noon)**

Remove the mating plate from the 4°C fridge.

Take it out of the ziplock bag, and put it near the fly workstation to warm up a little.

Wait about 5 minutes.

Next remove the FEEDING PLATE (containing fed-females) from the 25°C incubator

Invert the plate so that the perfusion-friendly lid is now on the bottom.

Place the CO_2_ gas pad right next to the hole on the side of perfusion-friendly lid.

As the flies fall asleep, they will drop to the (empty) bottom of the wells, and not get stuck.

While the treated female flies are falling asleep:

Use a second gas pad to anestheize the *w*Ri-infected males.

Separate the males into 24 sets of 10 flies (this should take about 5-10 min).

Remove the FEEDING plate to open/expose the perfusion-friendly lid and sleeping females

Use a funnel to quickly add 10 males to each well of the plate.

Then carefully place the MATING PLATE on top of the perfusion-friendly lid to close the plate.

Place strips of bench tape along 2 long sides of plate to secure the lid onto food section

*Make sure that alignment is good so flies can’t crawl from one well to another*

Wait until all the flies are fully awake before flipping plate over (*prevents flies from getting stuck*)

Place the plate in the 25°C fly incubator, using a middle shelf for best moisture control.

Avoid putting plate on bottom shelf of the incubator, as that seems to drive condensation.

Let the plate incubate for 8 hours.

After 8hr mating has concluded:

Remove MATING PLATE (with males/females, now mated) from the 25°C incubator.

Invert the plate so that the perfusion-friendly lid is now on the bottom.

Place the CO2 gas pad right next to the hole on the side of perfusion-friendly lid.

As the flies fall asleep, they will drop to the (empty) bottom of the wells, and not get stuck.

Remove the mating plate to open/expose the perfusion-friendly lid and sleeping females

Use dissection tweezers to remove every male from each well

Working quickly, be sure to recheck wells, confirm that all males were removed.

If flies start to wake up as you are picking the males out, place the gas pad over the open top of the perfusion-friendly lid to re-anesthetize the flies.

Then place the original FEEDING PLATE on top of the perfusion-friendly lid to close the plate.

Place strips of bench tape along 2 long sides of plate to secure the lid onto food section

*Make sure that alignment is good so flies can’t crawl from one well to another*

Wait until all the flies are fully awake before flipping plate over (*prevents flies from getting stuck*)

Place the plate in the 25°C fly incubator, using a middle shelf for best moisture control.

Avoid putting plate on bottom shelf of the incubator, as that seems to drive condensation.

Let the plate incubate for 24 hours.

**DAY 4 Start the egg collection**

Remove the EGG LAY PLATE from the 4°C fridge.

Take it out of the ziplock bag, and put it near the fly work station to warm up a little.

Wait about 5 minutes

Next, remove the fly FEEDING PLATE (containing fed/mated females) from the 25°C incubator.

Invert the plate so that the perfusion-friendly lid is now on the bottom.

Place the CO_2_ gas pad right next to the hole on the side of perfusion-friendly lid.

As the flies, fall asleep, they will drop to the (empty) bottom of the wells, and not get stuck.

Remove the feeding plate to open/expose the perfusion-friendly lid and sleeping females

Then carefully place the EGG LAY PLATE on top of the perfusion-friendly lid to close the plate.

Place strips of bench tape along 2 long sides of plate to secure the lid onto food section

*Make sure that alignment is good so flies can’t crawl from one well to another*

Wait until all the flies are fully awake before flipping plate over (*prevents flies from getting stuck*)

Place the plate in the 25°C fly incubator, onto a middle shelf for best moisture control

Avoid putting plate on bottom shelf of the incubator, as that seems to drive condensation.

Let the plate incubate for 24 hours.

**DAY 5 Stop egg lay, let eggs hatch.**

Remove the EGG LAY PLATE from the 25°C incubator.

Using a microscope view the egg lay through the perfusion-friendly lid.

Count approximately how many eggs are in each well.

If all contain approximately 50 eggs or more per well, the plate is ready to score.

If more eggs are needed, place females back on the plate and repeat after an hour or more.

Once desired egg lay is achieved:

Invert the plate so that the perfusion-friendly lid is now on the bottom.

Place the CO_2_ gas pad right next to the hole on the side of perfusion-friendly lid.

As the flies, fall asleep, they will drop to the (empty) bottom of the wells, and not get stuck.

Remove the EGG LAY PLATE, set aside. Make sure all females have been removed from plate. Either tap the perfusion friendly lid over the gas pad for collection, or dispose in the morgue.

Cover the EGG LAY PLATE by replacing the empty, perfusion-friendly lid back over the egg laying plate, or use a regular 24-well plate, flat-top lid.

Place the plate in the 25°C fly incubator

Avoid putting plate on bottom shelf of the incubator, as that seems to drive condensation.

Position the plate in middle of incubator for moisture control.

Let the plate incubate for 24-26 hours.

**DAY 6 Score egg hatch**

Remove the EGG LAY PLATE from the 25°C incubator.

Remove lid to open the egg lay plate.

Place the egg lay plate under the dissection microscope.

Calibrate the lighting, zoom and focus to maximize visibility of eggs/egg remnants

Scan the plate first to see if the wells are nicely seeded with eggs.

Minimum 50 eggs per well, for all wells, is required to qualify the plate as suitable for scoring.

If a quick scan indicates that the plate is likely worth scoring, then:

Use a needle to count/score how many eggs did/did not hatch.

Document the outcome separately for each well.

**Statistical analysis:**

When done, statistically compare the hatch rate between CI and CI+treatment conditions.

First do Shapiro-Wilk to test for normality of the data distribution.

Then use Levene’s test to assess homogeneity of variance.

Based upon those outcomes, you can choose appropriate statistical test for analyzing data

Shapiro-Wilk p < 0.05?	Levene p < 0.05?	Test to use for analyzing data

No (normal distrib)	No (equal variance)	T-test

No (normal distrib)	Yes (unequal variance)	Welch’s T-test

Yes (non-normal distrib)	No (equal variance)	Mann-Whitney

Yes (non-normal distrib)	Yes (unequal variance)	Indep. T-test w/ bootstrapping

We have typically found use of T-tests and Mann-Whitney covers most of the datasets we get.

On occasion we need Welch, and rarely need the Independent T-test with bootstrapping.

Note: Since the Independent T-test w/ bootstrapping is a more approximate test than the others:

We check, make sure that no experimental interpretation ever rests solely upon use of that test.

If all replicates of any given treatment needed that, we would plan another run of the experiment.

This analysis method is as per the publication: [*https://www.ncbi.nlm.nih.gov/pubmed/31481018*](https://www.ncbi.nlm.nih.gov/pubmed/31481018)

**Protocol: Fly rearing for CI assays**

**General maintenance**

Incubator (Invictus, Genessee Scientific, USA) was maintained at 25°C on a 12h light/dark cycle and used to house *Wolbachia* infected and uninfected (cured) *Drosophila simulans* stocks.

To obtain sufficient volumes of flies, 3-4 boxes of 16 bottles each per strain of *D. simulans* were maintained under standard conditions (see main text), transferring to fresh stock bottles approximately once per week.

Bottles produce a subsequent generation of flies within 10 to 12 days, thus experiments were scheduled accordingly.

Boxes were placed outside of the incubator, at room temperature, when pupal cases indicated the onset of the hatching pulse. Bottles were cleared of adult flies, double-checking to confirm the lack of all adult flies. This insured previously mated flies were absent from the population.

Depending on food consistency and to avoid the disturbance of pupa by liquid food when clearing or flipping bottles, a corner of ½ - 1 Kimwipe was submerged into the food using an ethanol disinfected spatula.

From the onset of hatch, bottles were used for approximately 5-7 days for experiments, collecting flies up to 8 hrs old or less to ensure virgin status, and separated into male or female populations. Bottles not used for experiments were directed to maintaining the stock population totals, with flies sourced from these bottles for up to 2 weeks for this purpose.

Stock quality was monitored regularly. Any bottle that accumulated mold or other microbial contamination indicated by reddish food discoloration, was immediately discarded. Fly infection status was also regularly reconfirmed by PCR.

**Bottle Flipping (cured, *w*Ri-infected and *w*Mel-infected)**

Approximately 2-3 days after initial seeding of bottles (or 3-4 days for *w*Mel), additional stock flies could be produced by flipping into unseeded bottles, or discarded if enough flies are available for setting up an appropriate quantity of plates.

Successive bottle ‘lineages’ were associated by marking with numbers (1-16 within a given box), which allowed the tracking of data and regular checks for cross-contamination between stocks.

PCR assessment of *Wolbachia* infection status was preformed intermittently to confirm presence or absence of the symbiont in stock lines (Christensen et al., 2016).

Bottle populations were limited to approximately 40 – 60 flies, to avoid overcrowding.
