## Additional File 3 for "A Role for Maternal Factors in Suppressing Cytoplasmic Incompatibility"

**Drug type**

Celastrol

NaBu

Cycloheximide

Teniposide

Bortezomib

and MG132

Trametinib

**DNA integrity**

Shown to cause DNA damage in some studies [1, 2]

Prevents DNA damage in certain conditions [3, 4]

__________________

Protects/prevents DNA damage [5, 6], in part by via antioxidant proteins [7] and in part via histone H4 acetylation [8, 9]

__________________

Prevents formation of single-stranded [10] and double-stranded [11] DNA breaks

__________________

Primarily creates single-stranded DNA breaks [12]

__________________

Inhibit DNA damage repair by acting on ATM kinase and other DNA repair pathway factors [13]

__________________

Inhibits DNA damage repair under certain circumstances [14], possibly by acting on ATM kinase [15]

_________________

**Cell cycle timing**

Induces cell cycle arrest [16, 17]

Down-regulates Cyclin D, which normally drives re-entry into G1 [18]

__________________

Inhibits cell cycle in a concentration-dependent manner [19], possibly by suppressing Cyclin D expression [20]

__________________

Slows the cell cycle, primarily in G1 and S phase [21, 22]

__________________

Inhibits cell cycle in a concentration-dependent manner [23]

__________________

Induce cell cycle arrest at G2/M [24]

__________________

MEK inhibitor, drives cell cycle arrest at G0 and G1 [25, 26]

__________________

**Protein turnover**

Inhibits proteasome activity [27]

Enhances effect of Bortezomib and MG132 [28, 29]

__________________

Stimulates proteasome activity [30]

__________________

Inhibits production of ubiquitin subunits [31]

__________________

**Induces proteasomal degradation of certain targets [32]

__________________

Directly inhibit proteasome activity [33]

__________________

**Induces proteasomal degradation of certain targets [34]

__________________

** *narrowly tested*
